# A Non-Synonymous SHARPIN Variant is Associated with Limbic Degeneration and Family History of Alzheimer’s Disease

**DOI:** 10.1101/196410

**Authors:** Sourena Soheili-Nezhad, Neda Jahanshad, Sebastian Guelfi, Reza Khosrowabadi, Andrew J. Saykin, Paul M. Thompson, Christian F. Beckmann, Emma Sprooten, Mojtaba Zarei, for the Alzheimer’s Disease Neuroimaging Initiative

## Abstract

Pharmacological progress, basic science and medical practice can benefit from objective biomarkers that assist in early diagnosis and prognostic stratification of diseases. In the field of Alzheimer’s disease (AD), the clinical presentation of early stage dementia may not fulfill any diagnostic criteria for years, and quantifying structural brain changes by magnetic resonance imaging (MRI) has shown promise in the discovery of sensitive biomarkers. Although hippocampal atrophy is often used as an in vivo measure of AD, data-driven neuroimaging has revealed complex patterns of regional brain vulnerability that may not perfectly map to anatomical boundaries. In addition to aiding diagnosis, decoding genetic influences on neuroimaging measures of the disease can enlighten molecular mechanisms of the underlying pathology in living patients and guide the therapeutic design.

Here, we aimed to extract a data-driven MRI feature of brain atrophy in AD by decomposing structural neuroimages using independent component analysis (ICA), a method for performing unbiased computational search in high dimensional data spaces. Our study of the AD Neuroimaging Initiative dataset (n=1,100 subjects) revealed a disease-vulnerable feature with a network-like topology, comprising amygdala, hippocampus, fornix and the inter-connecting white-matter tracts of the limbic system. Whole-genome sequencing identified a nonsynonymous variant (rs34173062) in SHARPIN, a gene coding for a synaptic protein, as a significant modifier of this new MRI feature (p=2.1×10^−10^). The risk variant was brought to replication in the UK Biobank dataset (n=8,428 subjects), where it was associated with reduced cortical thickness in areas co-localizing with those of the discovery sample (left entorhinal cortex p=0.002, right entorhinal cortex p=8.6×10^−4^; same direction), as well as with the history of AD in both parents (p=2.3×10^−6^; same direction).

In conclusion, our study shows that ICA can transform voxel-wise volumetric measures of the brain into a data-driven feature of neurodegeneration in AD. Structure of the limbic system, as the most vulnerable focus of brain atrophy in AD, is affected by genetic variability of SHARPIN. The elevated risk of dementia in carriers of the minor allele supports engagement of SHARPIN in the disease pathways, and its role in neurotransmitter receptor scaffolding and integrin signaling may inform on new molecular mechanisms of AD pathophysiology.

**Abbreviations:** Alzheimer’s disease (AD), genome-wide association study (GWAS), independent component analysis (ICA), mild cognitive impairment (MCI), medial temporal circuit (MTC), single-nucleotide polymorphism (SNP), tensor-based morphometry (TBM)

## Introduction

AD is the leading cause of death in the elderly for which no disease-modifying treatment yet exists, with over 500,000 annual mortalities attributed to this disorder in the United States^1^. Due to complex disease mechanisms and molecular pathophysiology, all drug candidates of AD have failed to slow the progression of dementia in clinical trials. Early diagnosis of AD for possible intervention is also difficult, since the mild and heterogeneous presentation of early stage dementia reduces accuracy of any diagnostic criteria, and sensitive biomarkers do not yet exist to guide the clinical diagnoses.

A number of structural MRI measures have been evaluated as in vivo features of AD, to be used to aid disease diagnosis and monitoring trajectories. These include loss of hippocampal volume^2^, reduced thickness of the entorhinal cortex^3^ and increased volume of lateral ventricles^4^. Most of these imaging features have been developed based on a priori assumptions of the boundaries of known anatomical structures and involvement of these structures in patients. However, the sensitivity of a priori driven approaches that single out a particular brain region such as the hippocampus is limited by heterogeneous disease mechanisms that translate to different patterns of brain degeneration in different patients^5^. The region of interest studies also disregard important facts such as spatial continuity of brain areas^6^ and the anatomical covariance of brain networks^7^. Exploratory methods can circumvent this problem by tracking the disease impact in a hypothesis-free manner and providing a data-driven picture of the affected brain.

AD is highly heritable at h2 = 58-79%^8^. However, APOE4, which is the strongest genetic risk factor for sporadic late-onset AD, explains only a quarter of this genetic variance, and the novel risk variants discovered by genome-wide association studies (GWAS) explain an even smaller proportion of AD heritability^9^. These studies commonly rely on diagnostic instruments for the dichotomous definition of the disease versus the healthy state. However, diagnostic instruments are originally aimed at guiding decisions in the clinic and capture mostly the terminal events of the disease pathway. For better monitoring the underlying pathology, in vivo neuroimaging phenotypes comprise one set of promising tools, especially when the mild clinical presentation of early stage AD does not fulfill any diagnostic criteria.

Here, we used independent component analysis (ICA) of T1-weighted brain MRI data in a longitudinal cohort of elderly subjects to arrive at a data-driven marker of brain degeneration in AD. We next performed a genome-wide search for genetic variants associated with structural integrity of this brain covariance feature. Finally, we replicated the genome-wide association results in an independent sample seeking association of the top-hit variant with brain structure and the familial risk for AD.

## Results

### Independent component analysis reveals structural covariance in diverse brain regions

The ADNI subjects included 1,100 individuals (age: 74.5±7.1y) with baseline T1-weighted brain MRI scans. Cognitive status of participants included cognitively normal, mild cognitive impairment (MCI) and probable AD based on NINCDS/ADRDA criteria (Table 1). Longitudinal MRI scans were available in a subpopulation of 1,039 subjects who underwent follow-up imaging at 1.07±0.08 year (n=1,009 subjects) and/or 2.07±0.11 year intervals (n=883 subjects).

**Table 1.** Study population.

|  | CN | MCI | AD | Total |
| --- | --- | --- | --- | --- |
| Cross-sectional | 383 | 456 | 361 | 1,100 (491 women) |
| Longitudinal | 269 | 422 | 348 | 1,039 (457 women) |
| CN: Cognitively normal; MCI: Mild cognitive impairment; AD: Alzheimer's disease |  |  |  |  |

Jacobian determinant maps were created from structural MRI volumes using the Tensor-Based Morphometry (TBM) method^10^. These 3D fields encode brain volume differences in each of the study subjects in relation to a common average brain template constructed by deformable registration^11^ of all subjects’ brains (Fig. S1 in the Supplementary Appendix). For those subjects who underwent serial MRI scans at one or two years after the baseline scan, subtle anatomical changes across inter-scan intervals were also quantified by symmetric alignment of baseline and follow-up MRIs, producing longitudinal versions of the Jacobian maps that encode brain atrophy in course of disease progression11 (Fig. 1).

**Figure 1.**
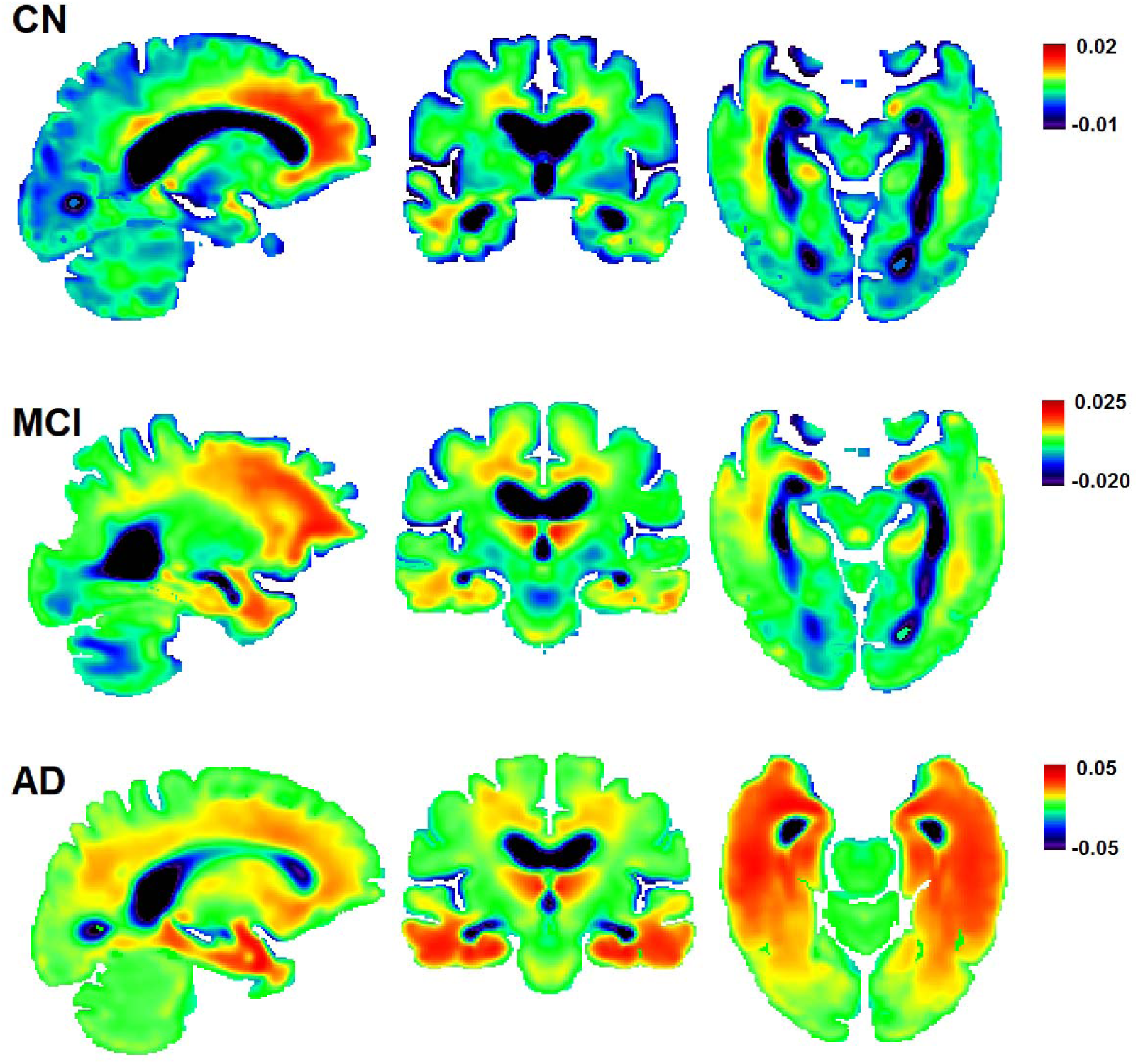
Tensor-based morphometry. Longitudinal Jacobian maps show the annual rate of brain atrophy in study diagnosis groups. CN: cognitively normal; MCI: mild cognitive impairment; AD: Alzheimer’s disease.

We then extracted independent features in brain structure by performing probabilistic ICA^12^ at various dimensions on the Jacobian maps of the whole study population. Cross-sectional (n=1,100 subjects) and longitudinal Jacobian determinant maps (n=1,039 subjects) were decomposed into a total of 1,348 spatially independent components across 1,068,867 brain voxels. The extracted components represent brain regions with structural covariance across the study population, such as areas that tend to lose volume due to a common underlying pathology (Fig. 2).

**Figure 2.**
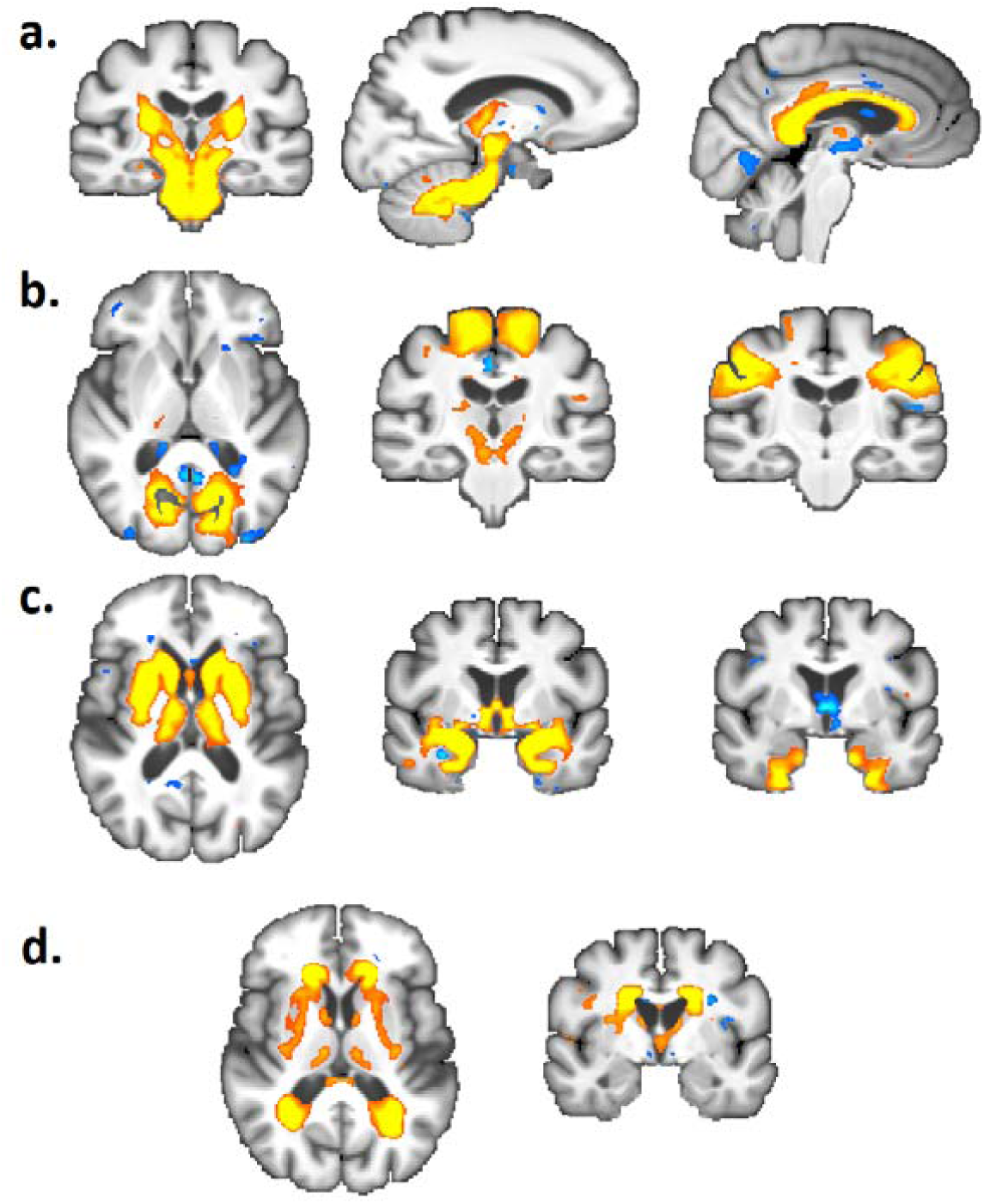
Independent components of structural brain variation. Examples of independent components decomposed in cross-sectional brain morphometry are shown. Components spanned various brain areas, including white-matter voxels in brainstem, cerebellum and corpus callosum (a), cortical gyri and sulci in occipital and frontal lobes (b), and subcortical grey matter including thalamus, striatum and hippocampus (c). Other features such as periventricular white matter lesions are also captured as independent spatial components (d).

To find out which of the 1,348 brain components were related to AD, they were used as predictor variables in regularized regression models^13^ to discriminate AD patients from the cognitively normal group. AD patients could be classified by reduced volume of 26 brain components in cross-sectional MRI, or 30 components in longitudinal MRI, with respective classification accuracies of 86% and 87% as assessed by leave-one-out cross validation (Fig. 3 and S3; for details, see methods section in the Supplementary Appendix).

**Figure 3.**
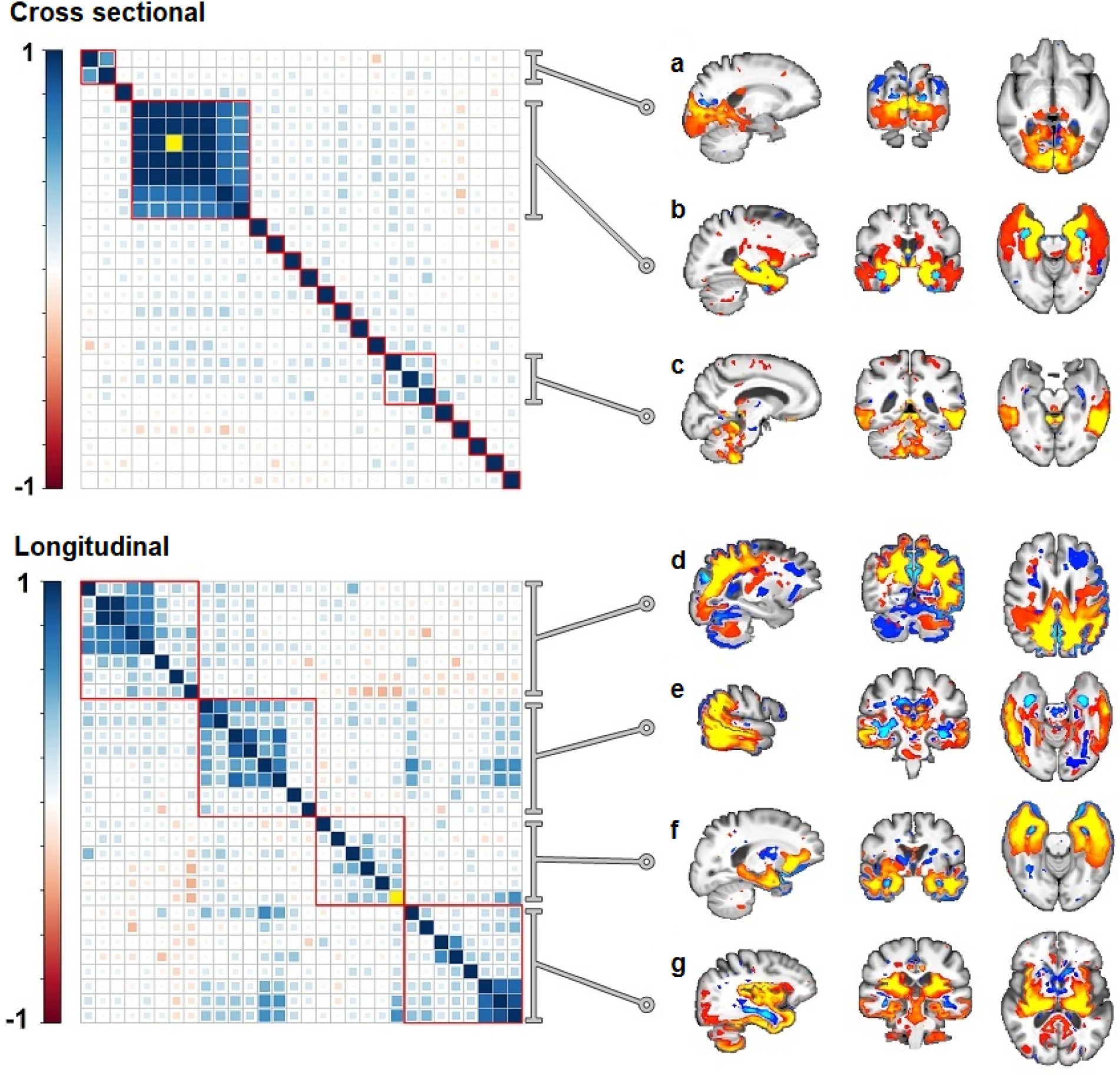
Brain components related to Alzheimer’s disease. Left: Cross-correlation matrices of brain components that discriminated AD patients from cognitively normal subjects were constructed. Hierarchical clustering was then applied to these matrices to group similar components together. Right: cluster-wise sum of the z-score maps of the components are shown (red-yellow: atrophy, blue: expansion; thresholded via mixture modeling^12^). The strongest AD discriminator in both analyses was a component referred to as the medial temporal circuit (MTC) in this paper, which is plotted as the yellow diagonal element in both matrices. This component (see volume rendered Video S1 in the Supplementary Appendix) was the “hot” focus of brain atrophy in a cluster of components mapping to temporal lobes (b).

The cross-sectional components predicting AD diagnosis clustered to: precuneus, occipital lobes and pulvinar (Fig. 3A); a robust cluster spanning hippocampi, amygdalae, parahippocampal gyri, fornix, mammillary bodies, and uncinate fasciculi (Fig. 3B); as well as lateral temporal lobes and the cerebellum (Fig. 3C). The AD-predicting components in longitudinal TBM clustered to: precuneus, cuneus and occipitoparietal lobes (Fig. 3D); thalamus, temporal lobes and its association areas (Figure 3E); medial temporal lobes, uncinate fasciculus and the orbitofrontal cortex (Fig. 3F); as well as insula, basal ganglia and cerebellum (Fig. 3G).

### Components of the limbic system show strong structural covariance in Alzheimer’s disease and suffer selective degeneration

Of the 1,348 ICA components decomposed, the most prominent imaging predictor of AD was a component that obtained the top odds-ratio rank in both cross-sectional and longitudinal analyses in discriminating AD from the cognitively-normal subjects. This brain component, which will be referred to as the medial temporal circuit (MTC) in this paper, was also a predictor of subjects with MCI, obtaining the top odds-ratio rank in discriminating MCI subjects from the cognitively normal group in cross-sectional brain morphometry (Fig. 4). MTC demonstrated a network-like topology in 3D and bilateral symmetry (see Video S1 in the Supplementary Appendix). The focus of brain atrophy in MTC localized to voxels of bilateral amygdalae, and further extended to hippocampi, entorhinal cortex, insula, mammillary bodies, and the fornical tracts.

**Figure 4.**
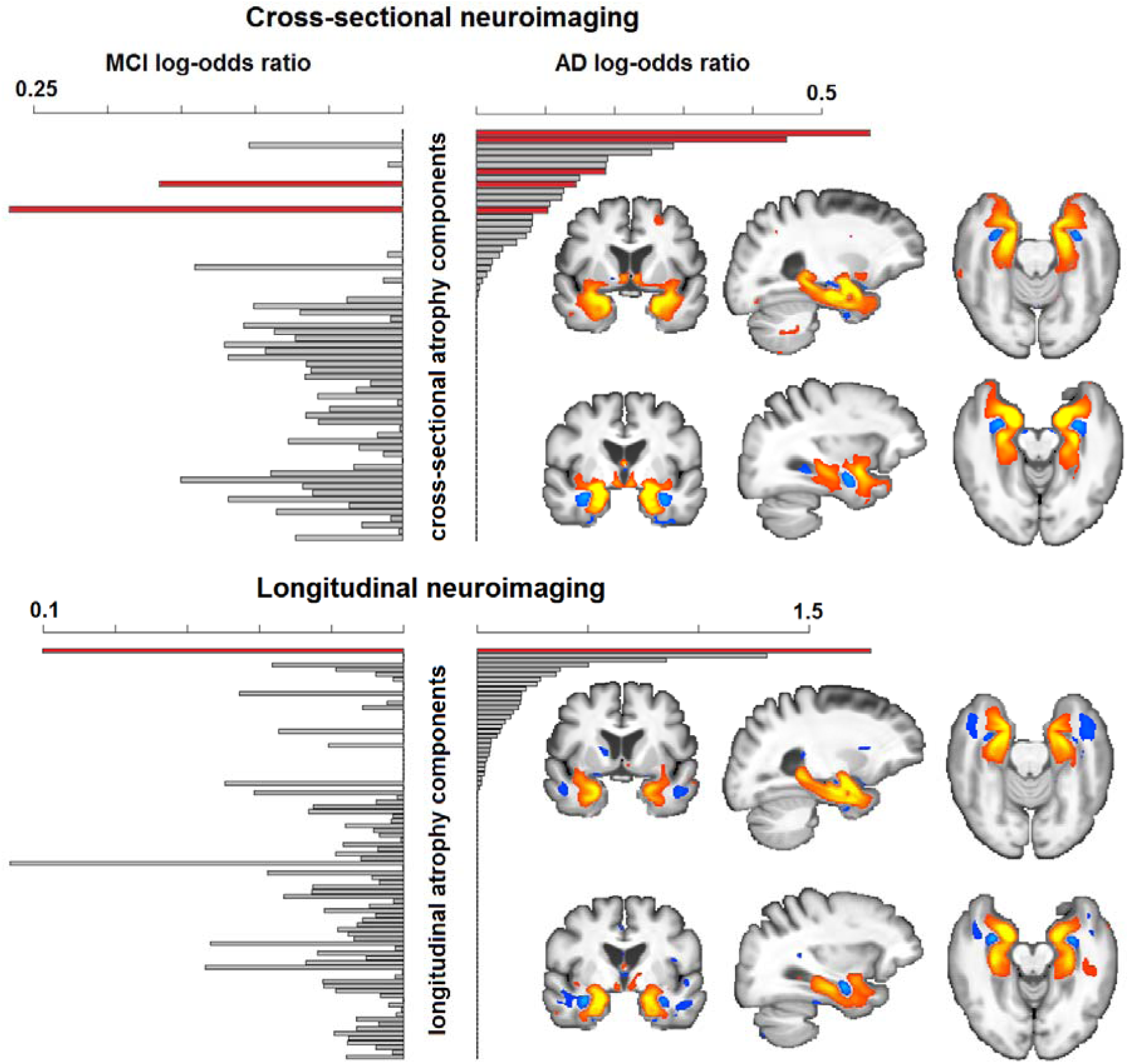
Contribution of brain components in predicting subjects’ cognitive status. Each bar represents a brain component that was able to distinguish MCI subjects (left plots) and/or AD patients (right plots) from the cognitively normal subjects in L1 regression models. Bar lengths encode log odds ratios reflecting the importance of each component in diagnosis classification. Both models in cross-sectional (top) and longitudinal neuroimaging (bottom) consistently identified the MTC component as the strongest predictor of AD and MCI (red bars). MTC was almost identically (r2>0.9) decomposed at five different ICA dimensions of cross-sectional analysis.

MTC was correlated with hippocampal volume (r=0.8), but compared to hippocampus, it could better predict MCI subjects from the cognitively normal group (Welch’s t=4.4: p=1.2×10^−5^ vs. t=3.2: p=0.002 for MTC and hippocampus, respectively). For completeness, MTC also showed higher statistical power than hippocampal volume for group-wide discrimination of the AD from the cognitively normal group, although subject to circularity (Welch’s t=16.8: p<10^−15^ vs. t=4.4: p=1.1×10^−5^ for MTC and hippocampus, respectively).

### Reduction of brain volume in MTC is correlated with cognitive decline

Cognitive performance of participants, as measured by ADAS-cog, was correlated with MTC at its baseline volume in AD (p=5.4×10^−8^) and MCI (p=9.6×10^−7^), adjusted for age and sex. Similarly, the atrophy rate in MTC showed significant associations with ADAS-cog in AD (p=0.04) and MCI subjects (p=0.002) adjusted for age and sex. Older age (p=2.6×10^−5^) and female sex (p=7.5×10^−5^) were associated with faster atrophy in this imaging feature. Compared to the cognitively normal group, the annual atrophy rate in the MTC was increased by 2.1-fold in MCI and 5.1-fold in AD patients.

### A common genetic variant modifies brain volume in MTC

Observation of MTC as a reproducible feature of brain degeneration in AD compelled us to study genetic determinants of its structural integrity across the study population. Whole-genome sequencing data was collected at an average depth of 30-40x in 808 participants, from which a total of 40,631,363 single-nucleotide polymorphisms (SNPs) were called^14^. We searched for common genetic variants (minor allele frequency > 0.01) correlated with baseline MTC volume in all study diagnosis groups with mean age (±SD) of 74.0±7.1 years, including 226 cognitively normal, 402 MCI and 180 AD subjects, and 363 of whom were women. A single missense variant in the SHARPIN gene, rs34173062, was genome-wide significant at p=2.1×10^−10^ (Fig. 5 and Fig. S4). Two other sub-genome-wide loci mapped to EPHA7 and FRMD4A introns (table 2).

**Table 2.** GWAS top hits.

| SNP | Chr | Position (hg19) | A1 | A2 (effect) | Frequency | $\beta$ | Locus | p-value |
| --- | --- | --- | --- | --- | --- | --- | --- | --- |
| rs56112946* | 3 | 197077194 | C | T | 0.03 | -0.15 | - | $3.1 \times 10^{-7}$ |
| rs3778470 | 6 | 94075684 | G | A | 0.12 | -0.15 | EPHA7 (intronic) | $1.5 \times 10^{-7}$ |
| rs149101079 | 8 | 69347181 | G | A | 0.04 | -0.15 | C8orf34 | $3.1 \times 10^{-7}$ |
| rs34173062 | 8 | 145158607 | G | A | 0.10 | -0.19 | SHARPIN (missense) | $2.1 \times 10^{-10}$ |
| rs28439901 | 10 | 14494941 | G | A | 0.08 | -0.15 | FRMD4A (intronic) | $3.2 \times 10^{-7}$ |

**Figure 5.**
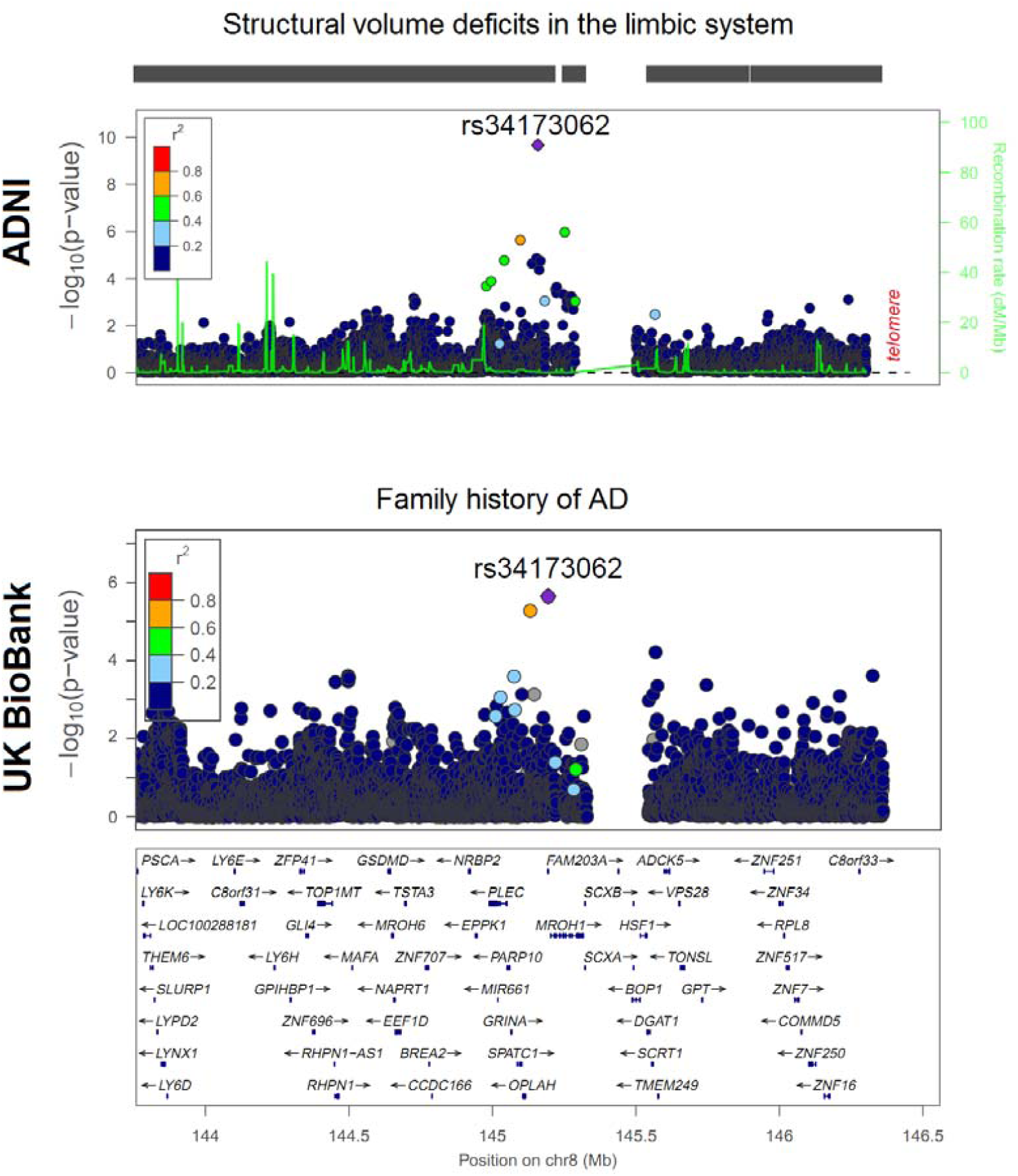
Regional association plots of the SHARPIN locus. Top: association of variants in the SHARPIN locus with structural deficits of the MTC feature in the ADNI cohort. Bottom: association of the same locus with parental history of AD in the UK biobank cohort.

We sought replication of the rs34173062 top-hit variant in the independent UK biobank cohort in which it was directly genotyped by a custom Axiom array^15^, showing a minor allele frequency of 0.07. In this cohort (n=8,428 subjects), rs34173062-A was correlated with reduced thickness of the entorhinal cortex in left (p=0.002) and right (p=8.6×10^−4^) hemispheres, with consistent effect directions (regression beta=- 0.1, Fig. 6). As the baseline age of the UK biobank subjects (49-69 years) is relatively young for clinical presentation of late-onset AD, we further searched for parental history of AD as a proxy to its heritable component^16^. The risk allele (rs34173062-A) was significantly correlated with increased subjects’ maternal (p=0.0012; n=308,780 subjects) and paternal (p=5.1×10^−4^; n=292,053 subjects) history of AD, which translates to a meta-analysis p-value of 2.3×10^−6^ in both parents with a consistent effect direction (Fig. 5: bottom).

**Figure 6.**
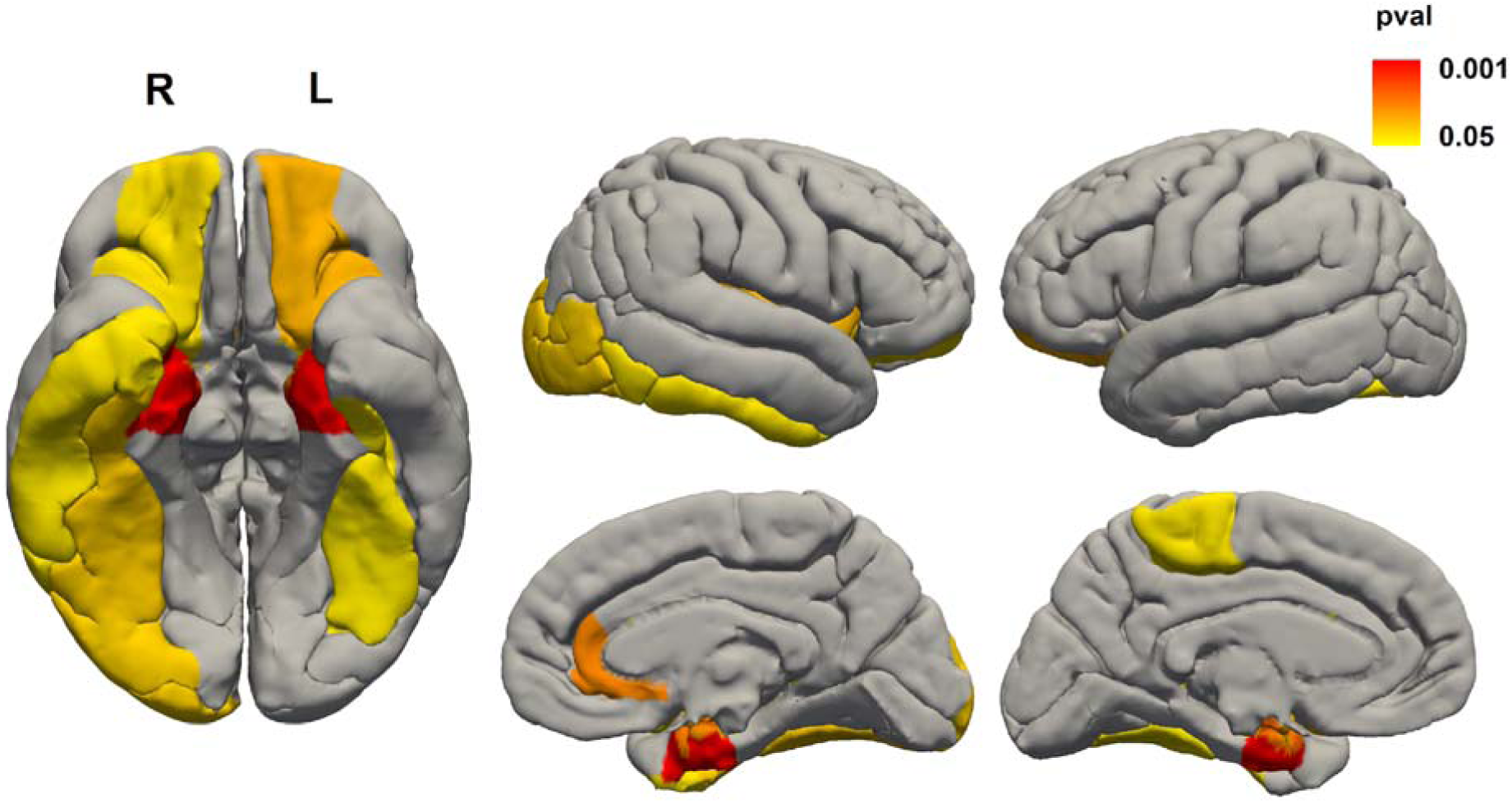
Association of rs34173062 with cortical thickness. Thickness of bilateral entorhinal cortices (red) was significantly associated with rs34173062 in the UK biobank cohort.

## Discussion

We observed strong covariance in structural deficits of the limbic system in AD. Considering the coherent role of this network in memory mechanisms, the voxel-wise covariance of brain volume loss in subcomponents of the limbic system may reflect network-level vulnerability of brain circuits to a common underlying pathology. This finding may be due to special cellular and proteomic compositions driving the observed gradient of brain volume loss in the ICA probability maps. Specifically, voxel-wise probability maps show that amygdalae are the most vulnerable structures of the limbic system in AD, with deficits further extending to bilateral hippocampi in a head-to-tail direction, and then to white matter connections including fimbriae and fornical tracts. In line with our findings, previous work has shown that healthy carriers of the APOE4 risk allele of AD suffer abnormal brain connectivity specifically in amygdalae and hippocampal heads, further suggesting that this region is affected early in the disease trajectory^17^.

In our whole-genome sequencing analysis of the MRI phenotype, we identified a nonsynonymous variant in the SHARPIN gene as a modifier of the medial temporal lobe structure in both ADNI and a relatively younger community-based cohort. This variant substitutes Phenylalanine for Serine in the N-terminal domain of the SHARPIN protein, a site responsible for its dimerization with potential roles in scaffolding due to adopting a pleckstrin homology (PH) superfold^18^. Association of this variant with the history of AD in both parents, in an independent cohort, verifies its contribution to dementia heritability, since each parent shares half of their genomic constitution with the child. Two recent articles have implicated SHARPIN in AD, with a rare variant (minor allele frequency: 2×10^−4^) in this gene found to increase the risk of AD to almost six fold, an effect larger than that of the TREM2 variant^19,20^. SHARPIN modulates recruitment of Kindlin-2, which is encoded by another AD risk gene, to the β1-integrin receptor, affecting signaling through the biological adhesion pathway^21^. As a component of the postsynaptic SHANK scaffold, SHARPIN also links the glutamate neurotransmitter receptors to the synaptic actin cytoskeleton^22^. Based on these observations, we suspect that SHARPIN may affect postsynaptic adhesion and scaffolding of the transmitter receptors with implications in synaptic stability. Our results add to the previous evidence by showing for the first time that common genetic variability in SHARPIN affects the structure of the vulnerable brain areas in two large cohorts, and increases the future risk of AD development as assessed by a parental by-proxy phenotype. Experimental studies in animals and cell cultures will be needed to determine which cellular pathways are responsible for the effect of SHARPIN on brain morphology and AD predisposition.

In conclusion, using linear separation of brain morphometry maps, we identify early limbic degeneration in AD as a new MRI feature, and report a genetic risk variant in SHARPIN associated with structural deficits of this vulnerable brain phenotype and elevated risk of AD.

## Author contributions

S.S-N., E.S. and M.Z. conceived and designed the study and wrote the manuscript. SS-N. analyzed the data. N.G., S.G, R.K., A.S., P.T and C.F.B. provided revisions to scientific content of the manuscript.

## Acknowledgements

Authors are thankful to Professor John Hardy and Professor Charles DeCarli for their useful comments. This work was supported in part by Radboudumc Hypatia Grant (R0003664 to E.S. and S.S-N.), NIH grants R01 AG059871 (N.J.) and R56 AG058854 (P.M.T), and research related grant support from Biogen PO 969323 (N.J. and P.M.T). Data collection and sharing for this project was funded by the Alzheimer’s Disease Neuroimaging Initiative (ADNI) (National Institutes of Health Grant U01 AG024904) and DOD ADNI (Department of Defense award number W81XWH-12-2-0012). ADNI is funded by the National Institute on Aging, the National Institute of Biomedical Imaging and Bioengineering, and through generous contributions from the following: AbbVie, Alzheimer’s Association; Alzheimer’s Drug Discovery Foundation; Araclon Biotech; BioClinica, Inc.; Biogen; Bristol-Myers Squibb Company; CereSpir, Inc.; Eisai Inc.; Elan Pharmaceuticals, Inc.; Eli Lilly and Company; EuroImmun; F. Hoffmann-La Roche Ltd. And its affiliated company Genentech, Inc.; Fujirebio; GE Healthcare; IXICO Ltd.; Janssen Alzheimer Immunotherapy Research & Development, LLC.; Johnson & Johnson Pharmaceutical Research & Development LLC.; Lumosity; Lundbeck; Merck & Co., Inc.; MesoScale Diagnostics, LLC.; NeuroRx Research; Neurotrack Technologies; Novartis Pharmaceuticals Corporation; Pfizer Inc.; Piramal Imaging; Servier; Takeda Pharmaceutical Company; and Transition Therapeutics. The Canadian Institutes of Health Research is providing funds to support ADNI clinical sites in Canada. Private sector contributions are facilitated by the Foundation for the National Institutes of Health (www.fnih.org). The grantee organization is the Northern California Institute for Research and Education, and the study is coordinated by the Alzheimer’s Disease Cooperative Study at the University of California, San Diego. ADNI data are disseminated by the Laboratory for Neuro Imaging at the University of Southern California.

## Supplementary methods

### ADNI participants

Imaging, whole-genome sequencing and clinical data used in preparation of this manuscript were obtained from the ADNI database (adni.loni.usc.edu). ADNI is a multi-center initiative led by principal investigator Michael W. Weiner, MD, VA Medical Center and University of California, San Francisco, and enrolls subjects with normal cognition, MCI and AD. Normal individuals have MMSE scores between 24 and 30, CDR of zero, non-depressed, non-demented and with no sign of cognitive impairment. Mild cognitive impairment subjects have MMSE scores between 24 and 30, memory complaint with objective memory loss measured by education-adjusted scores on Wechsler Memory Scale-Revised (WMS-R) logical memory II, a CDR of 0.5 and absence of significant impairment in other cognitive domains and absence of dementia. Alzheimer’s disease cohort had MMSE score between 20 and 26, CDR of 0.5 or 1.0 and fulfillment of NINCDS/ADRDA criteria for probable Alzheimer’s disease.

### MRI preprocessing and construction of common study template

Structural MRI was performed using GE, Philips or Siemens scanners. Structural T1-weigthed MRI volumes of the ADNI subjects were preprocessed for gradient non-uniformity and intensity inhomogeneity by the ADNI imaging core using gradient unwrapping, B1 field and N3 bias correction algorithms^23^. We registered all MRI volumes to construct a study brain template in four linear and four non-linear iterations using SyN^24^.

### Longitudinal atrophy estimation

The pre-processed T1-weighted volumes of subjects who underwent serial MRI scans were used to assess structural brain changes over time. FLIRT^25^ with 12 degrees of freedom was used to linearly transform a dilated mask in the common space of the study template into each subject’s native MRI space and obtain “cleaned” volumes in all subjects, which included brain and adjacent skull. This common stripping was performed to ensure that the most informative structures would be conserved for driving registration of serial MRI timepoints^26^, and unwanted anatomical differences (e.g. soft tissue changes) would be eliminated.

Basic steps of our longitudinal tensor-based morphometry (TBM) workflow have been extensively validated before^27,28^. The cleaned T1-weighted volumes of each subject at timepoints and timepoints were separately registered to halfway spaces (t_1,2-half_, t_1,3-half_) using 9 degree-of-freedom FLIRT with normalized-correlation cost function and FSL *midtrans* utility. Halfway space registration prevents bias stemming from arbitrary selection of an MRI time point as source or target in registration^29^. Nine degree-of-freedom registration removes global brain volume factor and has been shown to be comparable to phantom-based correction in ADNI^30^. In the halfway space, differential MRI bias correction was performed on source and destination imaging volumes using *BtkDifferentialBiasCorrection*, part of Biomechanical toolkit^31^. The calculated bias-correcting fields were transformed to native MRI spaces. This round of halfway space registration was used to calculate an identical skull-striping mask using logical AND for similar cropping of source and destination MRI volumes. After obtaining bias-corrected and identically-masked images in native scan spaces, another iteration of halfway space registration was carried out using the same methods. Final transformation matrices were subsequently converted to ITK format using *c3d_affine_tool* (www.itksnap.org^32^), and used to initialize a non-linear registration of source and destination time points by SyN. Parameters of longitudinal SyN registration included histogram matching, a course-to-fine resolution scheme with 20×10 gradient descent iterations, gradient step size of 0.25, an update field variance of 3, a total field variance of 0.5, and a cross-correlation metric with a search radius of 4 voxels.

### Independent Component Analysis

In cross-sectional phase of the study, fifteen ICA runs at a broad range of dimensions, spanning 8, 12, 16, 20, 24, 28, 32, 40, 48, 56, 64, 72, 80, 88 and 96 were used to decompose Jacobian maps into independent sources. Each resulting ICA component is represented by a spatial map, which is a 3D imaging feature common across the study population, together with a mixing profile vector, which linearly reflects the subject-wise brain volume parameter in the corresponding 3D source.

### Validation of the atrophy rate estimates

Linear evolution of brain atrophy in the temporal domain with a non-significant intercept at time point zero is a sign of biological plausibility in longitudinal TBM^28^. To validate our TBM workflow, the ICA-derived atrophy estimates were linearly regressed against the inter-scan interval duration in study diagnosis groups. Regression intercepts at baseline time point were not significantly different from zero in the cognitively normal (p=0.9), MCI (p=0.2) or AD subjects (p=0.7). To identify the impact of interscan interval duration on atrophy rate estimates, annualized atrophy rate parameters across interval_1→2_ were regressed against interval_1→3_ to seek their linear association. All regression models in the study diagnostic groups showed non-significant intercepts, as well as slopes acceptably near unity (Normal:1.07, mild cognitive impairment:0.97, Alzheimer’s disease:1.07, Fig. S2), indicating that our workflow provides linear atrophy estimates across the two-year timespan of this study.

### Sparse logistic regression models for diagnosis discrimination

All of the 684 or 664 brain atrophy components in the cross-sectional and longitudinal study phases were used in sparse logistic regression models to discriminate AD or MCI subjects from the cognitively intact group. Increasing the L_1_ regularization parameter (lambda) in sparse regression *shrinks* the model by setting the parameter estimates of the less relevant components to zero, resulting in inclusion of fewer components in diagnosis prediction. Leave-one-out cross validation was used to assess accuracy of the regression model at each lambda value and to find the optimization point (Fig. S3).

### Genome-wide association studies

Whole-genome sequencing was performed by a non-CLIA laboratory at Illumina^33^. DNA samples were sequenced on Illumina HiSeq2000 systems using paired end read with length of 100 bp at 30-40x read depth. Short read data were mapped to Human Genome build 19 by the proprietary Illumina variant caller CASAVA and single-nucleotide polymorphisms (SNPs) were called, resulting in a total of 40,631,363 variants. We considered SNP calls passing a minimum phred quality score of 30 (>99% accuracy) and read depths between a minimum of 10 and maximum of threefold average chromosomal depth, minor allele frequency of > 0.01, Hardy-Weinberg disequilibrium p-value of >10^−6^ and SNP call rate of > 0.9. The sequence-called SNPs were validated with the microarray-based genotyping results showing 99.8 ± 0.1% concordance.

The first three axes of population stratification were extracted by Eigenstrat^34^. Genome-wide association study of the MRI phenotype was conducted by Plink^35^ correcting for several confound variables including age, gender, dummy variables encoding diagnosis groups (AD, MCI, cognitively normal), three principal axes of population structure, MRI pulse sequence (MP-RAGE vs. SPGR), and the APOE4 allele dosage.

## Supplementary Figures

**Figure S1:**
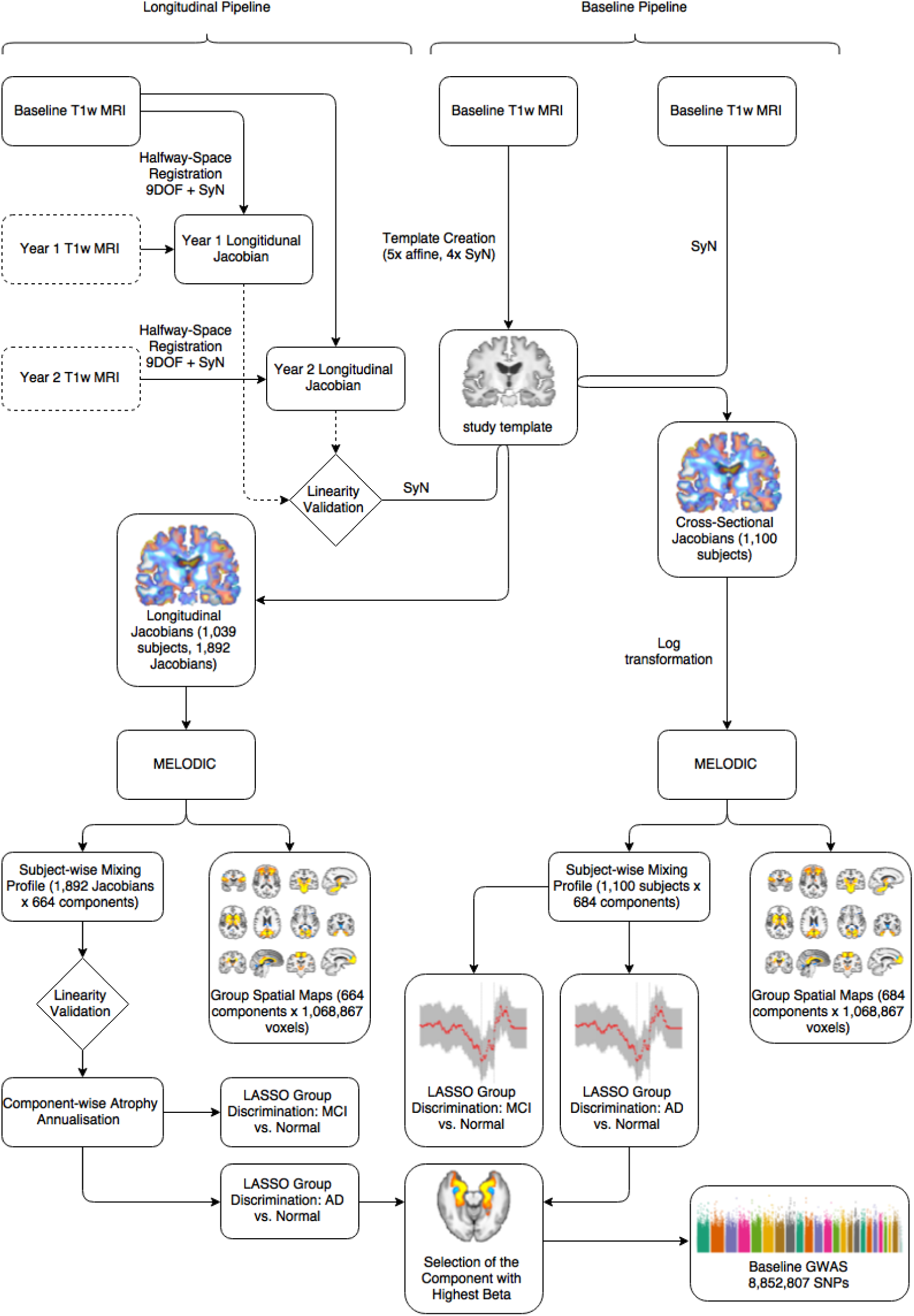
Outline of Study methods. T1-weighted MRIs were used to identify structural brain changes in cross-sectional and longitudinal studies. ICA decomposed 1,348 spatial sources of brain morphometry, among which the medial temporal circuit (MTL) was replicated as the top imaging predictor of AD and MCI and subsequently brought to GWAS.

**Figure S2:**
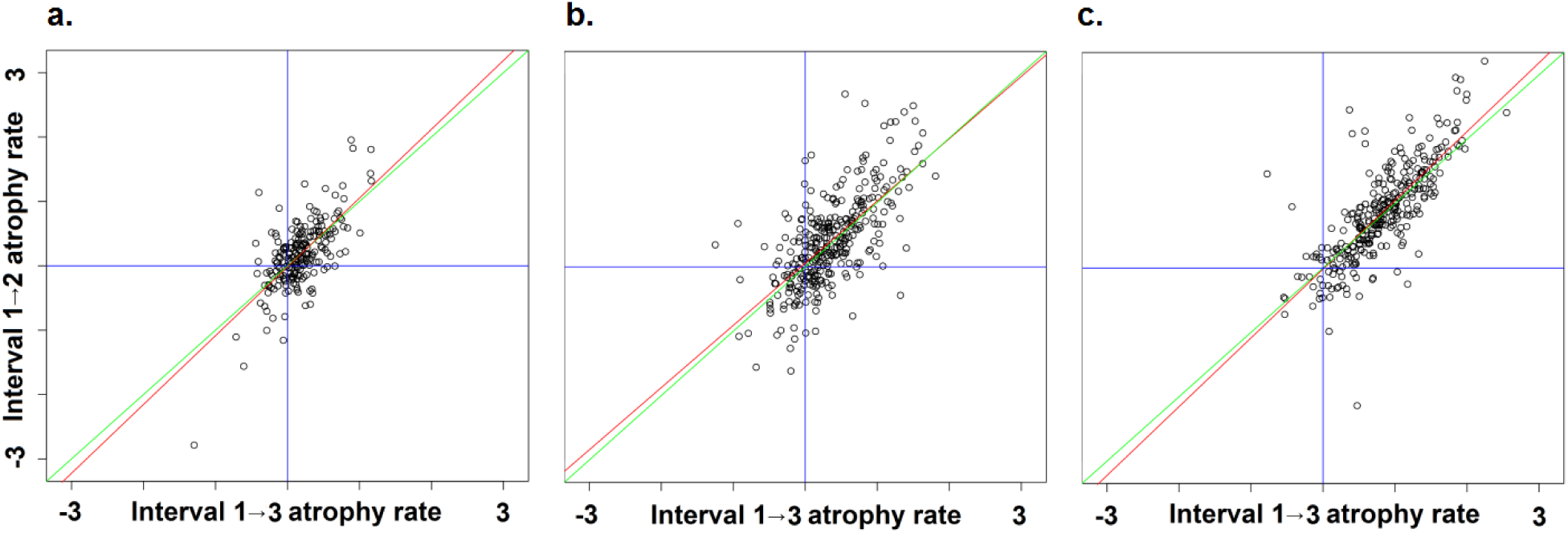
Colinearity of atrophy rates estimated by ICA. Annualized atrophy rate scatter plots of longitudinal MTL atrophy across timepoint_1→3_ (baseline and year-2) against timepoint_1→2_ (baseline and year-1) in cognitively normal (**plot A**), MCI (**plot B**) and AD (**plot C**) subjects. Regression lines (red) and the ground truth identity line (green) disclose minimal bias which was not statistically significant.

**Figure S3.**
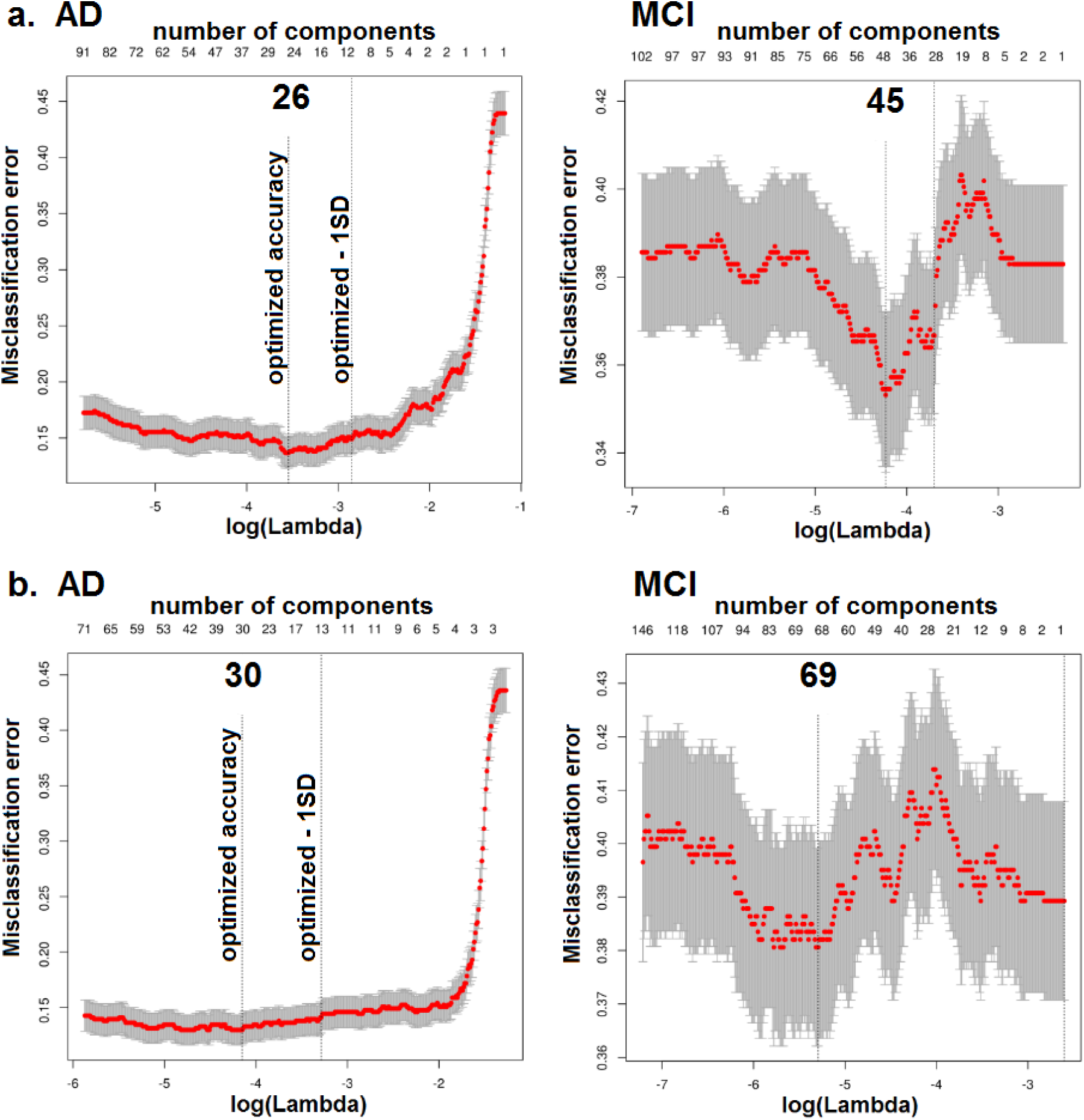
Sparse logistic regression models for diagnosis prediction. Leave-one-out cross-validation plots for discriminating AD or MCI subjects from the cognitively normal group using all ICA components. **a)** The increase in the λ parameter (x-axis: bottom) shrinks the L_1_ regression and results in inclusion of fewer predictor components in the diagnosis discrimination model (x-axis: top). Separate logistic regressions were used to discriminate AD (**left**) or MCI subjects (right) from the cognitively normal group. **b)** Similar regression models were used in longitudinal neuroimaging to discriminate AD (**left**) or MCI subjects (**right**) from the cognitively normal group.

**Figure S4.**
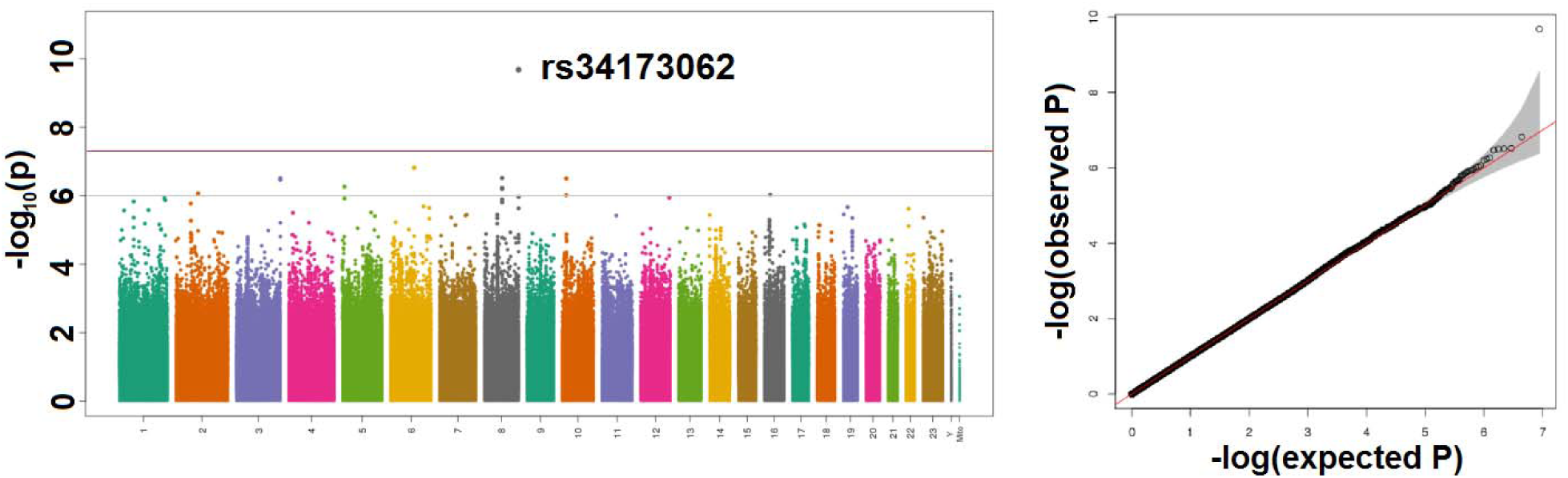
Manhattan and quantile-quantile plots of the medial temporal circuit association study. A common non-synonymous variant in SHARPIN was genome-wide significant.

## [Supplementary Video]

### 3D rendering of the medial temporal lobe component

The ICA z-score map of MTC is shown at incremental thresholds. The highest probability of brain atrophy maps to bilateral amygdalar voxels, and the effect further extends to the hippocampus, white-matter connections to fornix, mammillary bodies and other structures within the limbic system.

